# Structures of Langya virus fusion protein ectodomain in pre and post fusion conformation

**DOI:** 10.1101/2023.04.11.536446

**Authors:** Aaron J. May, Karunakar Reddy Pothula, Katarzyna Janowska, Priyamvada Acharya

## Abstract

Langya virus (LayV) is a paramyxovirus in the *Henipavirus* genus, closely related to the deadly Nipah and Hendra viruses, that was identified in August 2022 through disease surveillance following animal exposure in eastern China. Paramyxoviruses present two glycoproteins on their surface, known as attachment and fusion proteins, that mediate entry into cells and constitute the primary antigenic targets for immune response. Here, we determine cryo-EM structures of the uncleaved LayV fusion protein (F) ectodomain in pre- and post-fusion conformations. The LayV-F protein exhibits pre- and post-fusion architectures that, despite being highly conserved across paramyxoviruses, show differences in their surface properties, in particular at the apex of the prefusion trimer, that may contribute to antigenic variability. While dramatic conformational changes were visualized between the pre- and post-fusion forms of the LayV-F protein, several domains remained invariant, held together by highly conserved disulfides. The LayV-F fusion peptide is buried within a highly conserved, hydrophobic, interprotomer pocket in the pre-fusion state and is notably less flexible than the rest of the protein, highlighting its “spring-loaded” state and suggesting that the mechanism of pre-to-post transition must involve perturbations to the pocket and release of the fusion peptide. Together, these results offer a structural basis for how the Langya virus fusion protein compares to its Henipavirus relatives and propose a mechanism for the initial step of pre- to post-fusion conversion that may apply more broadly to paramyxoviruses.

**Importance:** The Henipavirus genus is quickly expanding into new animal hosts and geographic locations. This study compares the structure and antigenicity of the Langya virus fusion protein to other henipaviruses, which has important vaccine or therapeutic development implications. Furthermore, the study proposes a new mechanism to explain the early steps of the fusion initiation process that can be more broadly applied to the *Paramyxoviridae* family.

## Introduction

Langya virus (LayV) is a newly identified member of the *Henipavirus* genus, detected after surveillance of disease following animal exposure in eastern China (1). Over the course of several years, LayV infections were identified in 35 individuals, 26 of whom were infected solely with LayV. In these 26 individuals, common symptoms included fever, fatigue, cough, nausea, and headaches. The genome organization of LayV is identical to that of other henipaviruses, including the better known and highly virulent Nipah and Hendra viruses. Based on phylogenetic analysis, LayV is most closely related to Mojiang virus (MojV), which was discovered in rats in southern China (2), and Gamak (GAKV) and Daeryong (DARV) viruses, which were detected in shrews in the Republic of Korea (3).

The henipaviruses infect a range of animals with Nipah and Hendra viruses having their natural reservoir in fruit bats (4). However, several additional *Henipavirus* species have been discovered in recent years with differing animal reservoirs. A survey of both domestic and wild animals revealed a low level of seropositivity for LayV in goats and dogs (2% and 5% respectively) and the highest positivity rate (52.1%) in shrews (*Crocidura lasiura* specifically), indicating that shrews may be the natural reservoir for LayV. The *Henipavirus* genus is a member of the *Paramyxoviridae* family (5), which includes both extremely infectious human pathogens, such as measles and mumps, and extremely deadly pathogens, such as Nipah and Hendra. This diversity of infectivity and virulence, as well as diversity of animal reservoirs are risk factors that necessitate paramyxovirus research. Accordingly, Nipah and Hendra virus are listed as priority pathogens by the WHO (6).

Henipaviruses present two surface glycoproteins known as the attachment and fusion proteins. These proteins work together to mediate viral entry into host cells and in Nipah virus both have been shown to be required for viral entry (7). As these are the sole virus surface-expressed proteins, they are also the primary targets of neutralizing antibodies against henipaviruses. The *Henipavirus* attachment protein has receptor-binding capability and, in Nipah and Hendra viruses, binds to Ephrin B2 and B3, which is found mostly in the brain and in endothelial cells in the heart and lungs (8, 9). The *Henipavirus* fusion (F) protein is a class-I fusion protein that has a metastable pre-fusion conformation which is displayed on virion surfaces prior to receptor engagement and a post-fusion conformation that is adopted after virus-cell fusion (10, 11). Proteolytic cleavage splits F into two subunits, F_1_ and F_2,_ still connected by disulfide linkages (10, 12), freeing a series of hydrophobic residues known as the fusion peptide to be inserted into the host cell membrane during this conformational conversion. This process anchors the virus to the host cell and allows the formation of a six-helix bundle in the F protein to bring the viral and host membranes together, facilitating fusion. Although it has many mechanistic features in common with other class-I fusion proteins, the F protein in paramyxoviruses does not appear to have any primary receptor-binding functionality. This role is fulfilled by a separate attachment glycoprotein, which is believed to undergo conformational changes during receptor binding which pass a triggering signal to F (13, 14). The structural basis for this process is poorly understood.

Here, we determine cryo-EM structures of the LayV fusion protein (LayV-F) in pre- and post-fusion conformations, both at 4.64Å resolution, with both structures obtained from the same cryo-EM dataset. The LayV-F ectodomain sequence as reported by Zhang *et al*. (2022) was used (1), replacing the transmembrane and cytoplasmic domains with purification tags and a trimerization domain. No prefusion stabilizing mutations were made, nor was the fusion protein cleaved or otherwise knowingly induced to convert to the post-fusion state. Therefore, the presence of both conformations suggests that the protein was in a state of spontaneous conversion during vitrification. We compare it to other known *Henipavirus* fusion protein structures and elucidate the structural basis for pre- to post-fusion conversion in both LayV and more broadly for *Henipavirus* fusion proteins. Our study demonstrates that the highly conserved paramyxovirus fusion protein architecture is utilized by LayV-F, identifies a region of variability among *Henipavirus* fusion proteins in an important antigenic site, and provides evidence for a mechanism to describe how *Henipavirus* fusion proteins are triggered to undergo conformational changes during the fusion process.

## Results

### Purification and Structural Determination of LayV-F

We purified the LayV-F ectodomain based on protocols previously developed for NiV-F, by expressing in 293F cells and purifying via StrepTactin (IBA lifesciences) affinity chromatography followed by size exclusion chromatography. Like the NiV-F protein, the LayV F ectodomain elutes with a distinct peak at ∼180 kDa. Higher molecular weight species, broader than the main peak, were also observed at ∼400 kDa and above (Supplemental Figure 2A). These could be the result of low-affinity interactions between individual trimers. SDS-PAGE analysis of these high molecular weight species in both NiV-F and LayV-F purifications has consistently produced identical bands corresponding to the ∼180 kDa species, consistent with the uncleaved F protein (Supplemental Figure 2E). Higher order multimers of Henipavirus F proteins with rings described as “hexamers of trimers” have been previously reported (15). While larger molecular weight species were detected during purification of LayV-F (Supplemental Figure 2A), they comprised a far lower proportion of total protein as compared to purifications of wild-type NiV-F and a prefusion-stabilized NiV-F mutant, NiVop08 (Supplemental Figure 2A) (16). The yield of LayV-F at ∼1.2mg/L was considerably higher than that of wild-type NiV-F at ∼250µg/L. The NiVop08 yield was vastly higher than the unstabilized NiV-F construct at ∼3mg/L, consistent with the general trend of prefusion-stabilized fusion protein constructs expressing more efficiently than their wild-type counterparts (17–20). Differential scanning fluorimetry (DSF) suggested reduced stability of LayV-F compared to NiV-F (Supplemental Figure 2B). The LayV-F DSF profile closely resembles that of wild-type NiV-F, but is left-shifted, indicating that the temperature-induced conformational changes or unfolding events are occurring at a lower temperature for LayV-F.

We determined the structure of the LayV-F ectodomain trimer using cryo-EM. Reference-free 2D classes revealed both pre- and post-fusion conformations of LayV-F in the cryo-EM dataset (Figure 1A-B). After several rounds of classification and sorting we obtained 4.64 Å reconstructions (calculated within cryoSparc according to the FSC 0.143 gold-standard criterion) of the pre-fusion and post-fusion forms of the LayV F protein from approximately 200,000 and 80,000 particles, respectively. Assessment of the three-dimensional FSC (3DFSC) plot for these maps indicated resolutions of 5.42 Å and 5.25 Å for the pre- and post-fusion conformations, respectively (21). The structures determined revealed pre- and post-fusion architectures that were similar to those determined for other paramyxoviruses (15, 22–26). The pre-fusion LayV F structure has a compact “clove-like” shape with a large internal cavity and spans ∼110 Å from the bottom of the stalk to the top of the apex. In comparison, the post-fusion structure is elongated, spanning ∼160 Å, with regions that formed the pre-fusion apex straightening into a central coiled-coil, ending with the trademark six-helix bundle of the post-fusion conformation.

**Figure 1:**
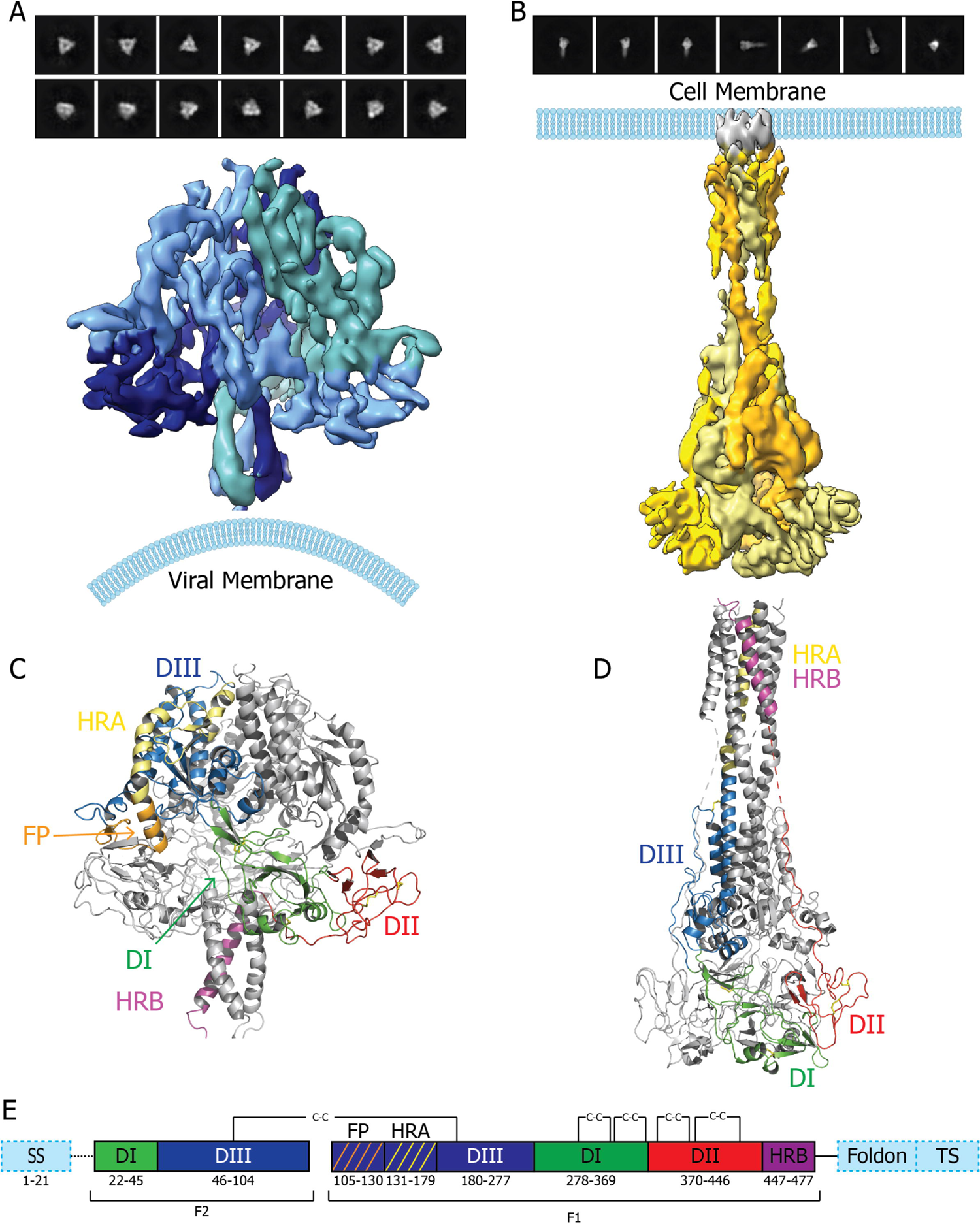
Structures of LayV-F in pre- and post-fusion conformation: **A-B.** Cryo-EM reconstructions of LayV-F in **A.** pre- and **B.** post-fusion conformation, colored by protomer. Map regions colored in different shades of blue (pre-fusion) and yellow (post-fusion) for each protomer. Representative reference-free 2D classes used for the reconstruction are displayed above each map. **C-D.** Cartoon representation of atomic models of LayV-F in **C.** pre-fusion and **D.** post-fusion conformation that were built into the maps shown in **A-B**. In each, one protomer is colored by domain, based on the color scheme in panel **E**, and the other two protomers colored gray. **E.** Sequence domain key for LayV-F. SS: Secretion signal, cleaved prior to purification. DI, DII, DIII: Domains I, II, and III. FP: Fusion Peptide. HRA, HRB: Heptad repeats A and B. Foldon: Trimerization domain. TS: Twin-Strep tag, used for purification with modified streptavidin resin.

The LayV F ectodomain includes a fusion peptide (FP) and two heptad repeat sequences that both contribute to three domains, previously described by Yin *et al*. for the parainfluenzavirus 5 (PIV5) F protein (Figure 1C-E) (26). Domain I (DI) is comprised of the most N-terminal residues after cleavage of the signal peptide, 22-45, and residues 278-369. Its central β-sheet connects Domain II (DII) to the apical Domain III (DIII). DII, from residues 370-446, makes significant contact with the cleavage site and fusion peptide of the neighboring protomer before leading to heptad repeat B (HRB), located at the most C-terminal end of the ectodomain. DIII, from residues 46-277, contains the cleavage site, fusion peptide, and heptad repeat A (HRA) (Figure 1C-E). Residues 93-140 from DIII and 441-450 from DII, which become linker regions leading to the repositioned heptad repeats now forming the six-helix bundle, are not resolved in the post-fusion structure (Figure 1E).

### Pre- to post-fusion Conversion Mechanism of LayV-F

Though the *Henipavirus* genus was established with the discovery of Hendra virus in 1994 (4, 27), the number of members of the genus has dramatically expanded in recent years with the discovery not only of Langya virus, but also Gamak and Daeryong virus, both identified in 2021 (3). Within the past three years, Henipaviruses have been identified in an expanded range of animal hosts and geographic locations. No longer contained to southeast Asia and Australia, new species have been discovered in Africa and South America, specifically Angavokely virus and Peixe-Boi virus, respectively (28, 29). Although there is very high structural similarity for fusion proteins across not only the *Henipavirus* genus, but within the larger *Paramyxoviridae* family, there tends to be low sequence identity between species, with LayV-F having ∼40% sequence identity with NiV-F, and ∼30% or less identity to other non-*Henipavirus* paramyxoviruses (Supplemental Figure 1). However, with more *Henipavirus* members identified, some conserved sequence features have become more evident. These include 10 highly conserved cysteines present in the ectodomain of paramyxovirus F proteins (Figure 2). Identifying the locations of these cysteines in the pre- and post-fusion LayV-F structures reveals their critical role in maintaining domain fold integrity during the dramatic conformational changes that occur during fusion.

**Figure 2:**
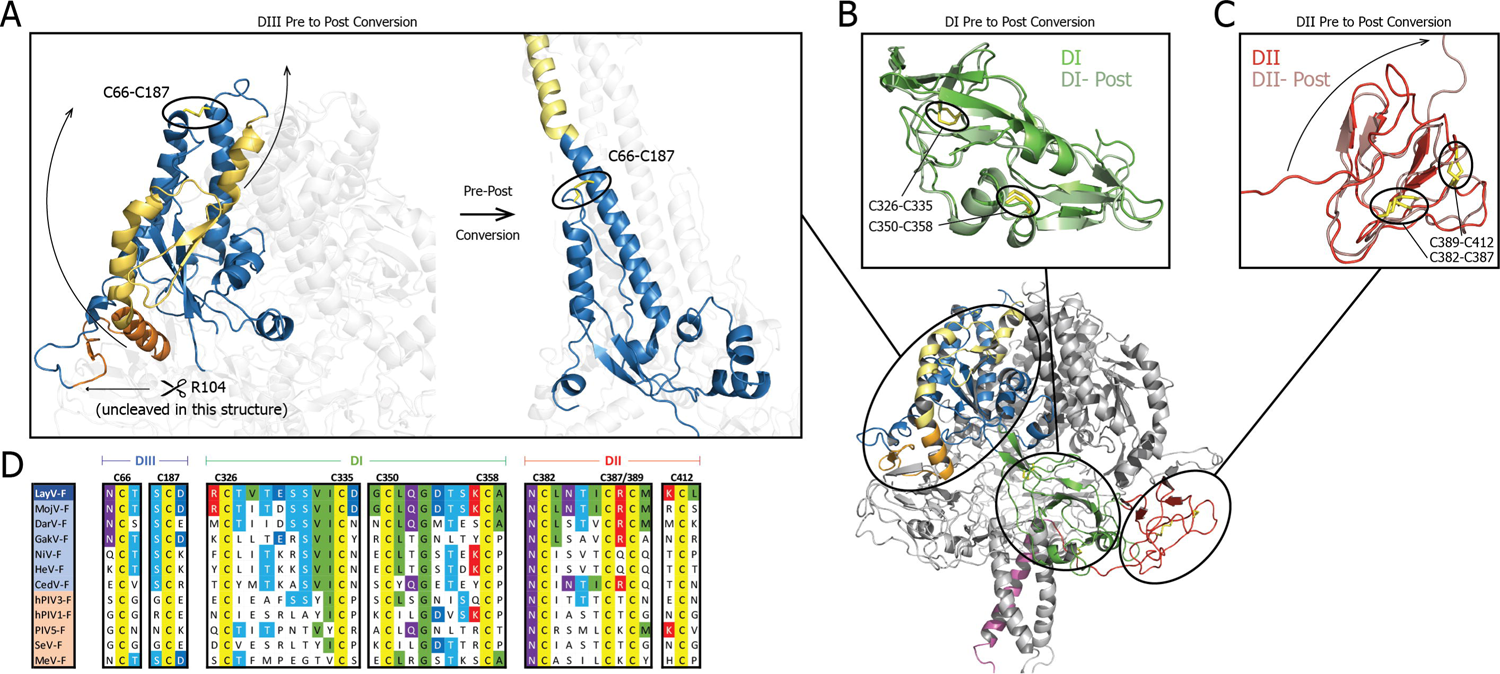
Role of conserved cysteines during conformational conversion. **A.** Zoomed-in view of the DIII domain, including the fusion peptide (orange) and HRA (yellow), shown for the (left) pre- and (right) post-fusion structures. Color scheme continued from Figure 1E. The cleavage site at R104 is identified, as well as the disulfide bond formed by C66 and C187 (yellow sticks). This disulfide bond is a hinge point around which the HRA straightens into the central coiled-coil in the post-fusion F structure. **B-C.** Superimposition of pre- and post-fusion conformation domains DI (**B**) and DII (**C**). Color scheme continued from figure 1E, with the post-fusion conformation colored in a lighter shade. Disulfide bonds are shown as yellow sticks and labelled. For DII, the movement of the HRB linker region is indicated by an arrow. **D.** Sequence alignment of paramyxovirus fusion proteins in the regions surrounding conserved cysteines, with cysteine residue number and domain labelled. Viruses colored blue are from the *Henipavirus* genus and colored tan for other paramyxoviruses. Residues are colored by type, per MView standard (50). Blue: Alcohol, Green: Hydrophobic, Dark Blue: Negative charge, Red: Positive charge, Purple: Polar, Yellow: Cysteine.

Despite the drastically different shapes of the pre- and post-fusion conformations of LayV-F resulting in a 51.6 Å Cα RMSD, superimposition of the DI and DII domains demonstrates that these domains remain largely invariant through the pre- to post-fusion transition of the F protein (Figures 2A-C). Both DI and DII domains have two internal pairs of disulfide bonds that help maintain the rigidity of these domains (Figure 2B-D). DI shows a pre- to post-fusion Cα RMSD of 2.16 Å, while DII undergoes more total movement, resulting in pre- to post-fusion Cα RMSD of 16.5 Å with the linker region from S423 to I446 that leads to HRB extending away from the domain as HRB moves to its post-fusion location (Figure 2C). Excluding these residues from the RMSD calculations, DII shifts only 2.3 Å. The difference in conformation of DIII from pre- to post-fusion is more dramatic. Based on sequence alignment, the F_1_-F_2_ cleavage site for LayV-F is expected to be immediately after the highly conserved residue R104 (Supplemental Figure 1), however cleavage did not occur in this construct (Supplemental Figure 2C-D). The largest positional shifts during the pre- to post-fusion transition occurred in the fusion peptide and HRA, which extend and continue the along the path of the apical helix, now forming the central coiled-coil (Figure 1A). These movements occur around a hinge point in DIII where the apical helix is connected to the rest of DIII by a disulfide bond, again demonstrating the role of the conserved cysteines in providing an anchor for large conformational changes in the F protein (Figure 2A).

### Fusion triggering Mechanism of *Henipavirus* Fusion proteins

Insertion of the hydrophobic fusion peptide (FP) into the host membrane is an essential step that leads to the fusion of the virus and host cell membranes. In the pre-fusion LayV-F, a substantial portion of the FP is buried within a pocket formed between two LayV-F protomers, requiring the FP to be released from this pocket during the pre-to-post fusion conformational transition (Figure 3). Hydrophobic residues lining the interprotomer pocket into which the FP is buried in the pre-fusion LayV-F include L51, L89, M92, L93, V96, V216, and I262 from DIII, A108, M110, G112, A114, L115, and G116 from the N-terminal end of the FP upstream of the helical segment from the same protomer, and residues from the adjacent protomer include M292 from DI and P369, F371, A372, L373, G376, V378, I420, L421, and I422 from DII (Figure 3D, Supplemental Figure 5).

**Figure 3:**
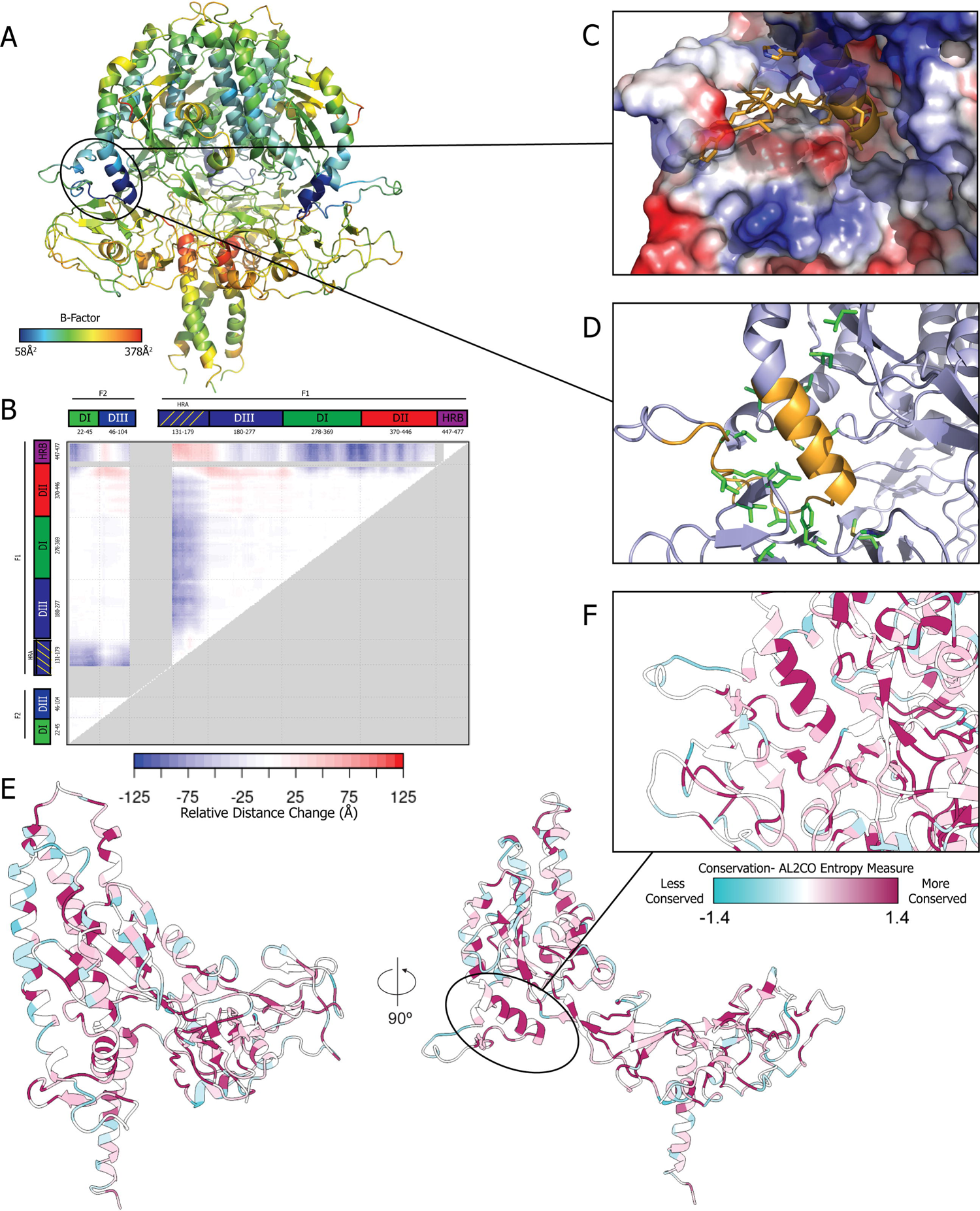
Fusion peptide burial and conservation. **A.** Structure of the pre-fusion LayV-F protein shown in cartoon representation and colored by B-factors. **B.** Difference Distance Matrix plot comparing relative residue positions between pre-fusion and post-fusion conformation. Sequence key from figure 1E included along each axis. Gray areas in the bottom right half of the plot are the result of sequences not present in the post-fusion model. **C.** Hydrophobic fusion peptide pocket in pre-fusion state, with fusion peptide depicted as orange cartoon, and surface electrostatics view of the pocket. **D.** Side-chain stick view of the fusion peptide pocket, with hydrophobic residues involved shown with side chains as green sticks. **E.** Cartoon representations of a single pre-fusion state protomer, colored by sequence conservation across *Henipavirus* fusion proteins. Color scale generated by ChimeraX default AL2CO entropy measure. **F.** Zoom in of the fusion peptide pocket, colored by sequence conservation as in H. The fusion peptide pocket is among the most strongly conserved region of the protein.

Consistent with the close packing of the FP within an interprotomer pocket in the pre-fusion F conformation, the modeled coordinates exhibit the lowest B-factors at the FP, while HRB, its DII linker, and surrounding DI region have the highest B-factors in the structure (Figure 3A). The HRA, HRB and FP undergo dramatic shifts in their positions during pre- to post-fusion conformational change of the LayV-F protein (Figure 3B). Although the FP is not resolved in the post-fusion state, based on its location it is expected to show a position shift similar to that of HRA. The burial of the pre-fusion FP and its interactions with the residues lining the interprotomer pocket suggest that the purpose of this pocket is to hold the FP in place to prevent premature conversion to the post-fusion form and therefore is the source of metastability of the pre-fusion LayV F structure. Consistent with a key role in pre-to post-fusion conformational dynamics, the FP and the surrounding pocket are among the most conserved regions in *Henipavirus* fusion proteins (Figure 3E-F). Release of the FP from this pocket is a requirement for the conformation transition of the F protein from the pre-fusion to post-fusion conformation and disruption of this pocket may be the mechanism by which the FP is released to undergo these changes.

### Structural Conservation and Antigenic Diversity of *Henipavirus* Fusion proteins

Despite low overall sequence conservation, the structural similarity of paramyxovirus fusion proteins is striking. Structures have previously been determined for the F proteins of two *Henipaviruses* in the pre-fusion state, NiV-F and HeV-F (15, 24). Despite having only 38% sequence identity with these viruses, a superimposition of LayV-F with these structures reveals a conserved architecture (Figure 4A), with Cα RMSDs of 2.64 Å and 3.00 Å for LayV-F to NiV-F and HeV-F, respectively, with NiV-F and HeV-F differing between each other by 0.94 Å. One of the few areas between these species where there is a notable shift is in HRA. In the pre-fusion conformation, the central helix in HRA is broken by a short, two-strand beta sheet with a loop that extends to the exterior of the protein. As compared to the homologous HRA loop in NiV/HeV-F, in LayV-F there is a shift of ∼9 Å in the direction of the fusion peptide (Figure 4B). The cryo-EM reconstruction in this region at an estimated local resolution of ∼5Å, (Supplemental Figure 3), is resolved well enough to allow unambiguous assignment of the main chain for this loop. It is unclear if this shift results in any mechanistic effect on the HRA region, although the loop resides within a site targeted by antibodies and shift in its position may alter antigenicity of the F protein. (Figure 4C).

**Figure 4:**
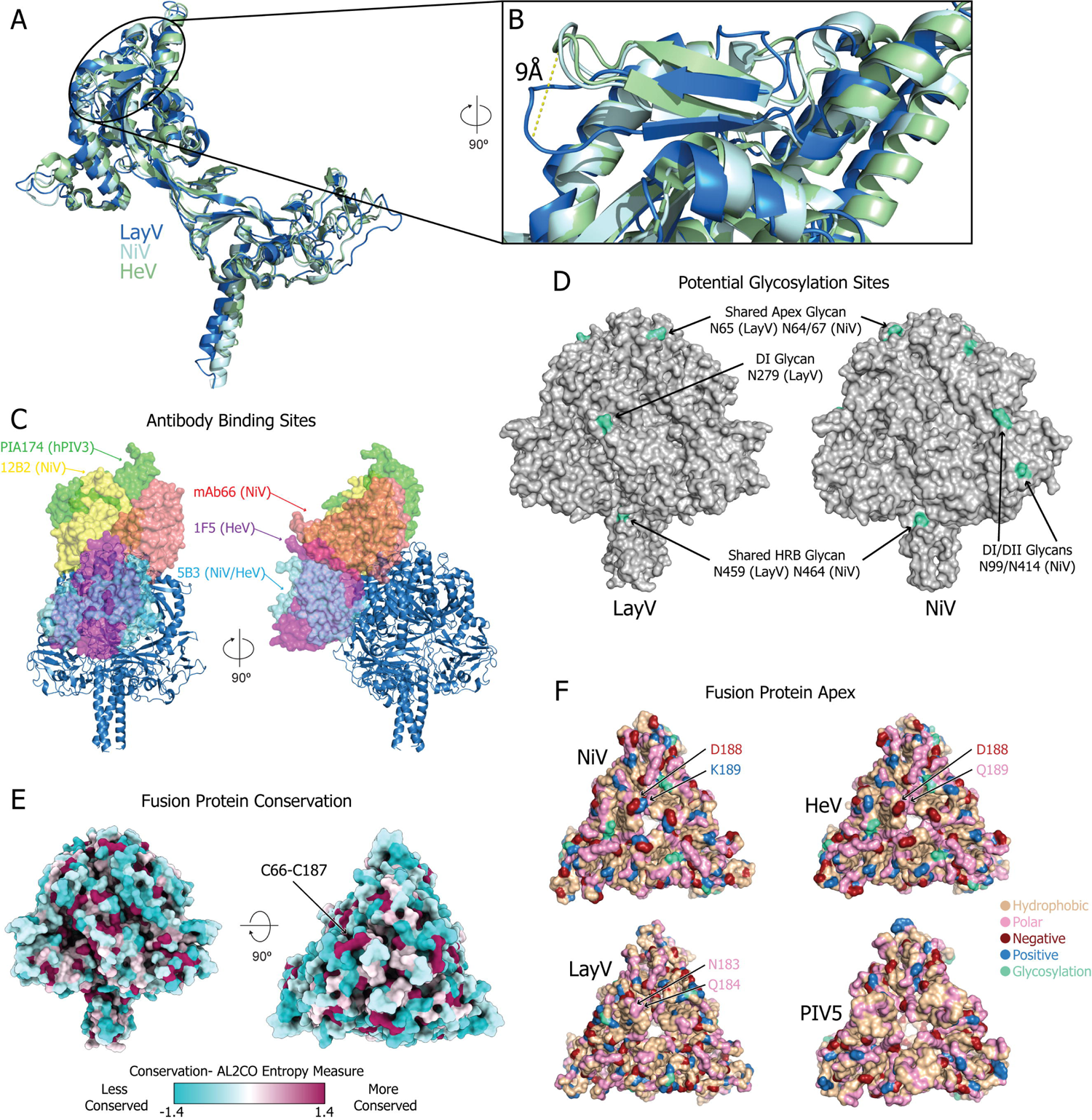
Structure and antigenicity comparison of Henipavirus fusion proteins. **A.** Superimposition of a single pre-fusion protomer of LayV-F (blue) NiV-F (cyan) and HeV-F (green), in cartoon representation. PDBs: LayV-F: 8FEJ, NiV-F: 5EVM, HeV-F: 5EJB (15, 24). **B.** Zoomed in view of a loop within the folded HRA, one of the few areas of significant structural difference between LayV-F and NiV/HeV-F. **C.** Binding site of known paramyxovirus F protein-neutralizing antibodies, labeled by the species targeted. NiV-F trimer from 12B2-bound structure shown as cartoon and antibody Fabs shown in surface view. Only one Fab per type shown, all expect PIA174 bind with 3:1 stoichiometry. PDBs: PIA174: 6MJZ, 12B2: 7KI4, mAb66: 6T3F, 1F5: 7KI6, 5B3: 6TYS (25, 30, 51). **D.** Surface view of LayV-F and NiV-F with glycosylation sites (residues meeting the N-X-S/T glycosylation sequence) colored teal. **E.** Surface representation of LayV-F colored by sequence conservation across *Henipavirus* fusion proteins. Color scale generated by ChimeraX default AL2CO entropy measure; magenta indicates high conservation and teal indicates high variability (52). **F.** Surface representation of paramyxovirus fusion proteins, showing only main chain atoms, colored by residue type and shown from top-down view. Tan: hydrophobic, Pink: polar, Red: negative, Blue: positive, teal: Potential glycosylation. PIV5 PBD: 4GIP (23).

An overlay of all experimentally determined structures of paramyxovirus-targeted antibodies highlights the antigenicity of the HRA loop site and its adjacent regions (Figure 4C, Supplemental Figure 5). Several antibodies target DIII in the pre-fusion F near the HRA region and likely interfere with pre- to post-fusion conversion. In addition to the HRA loop shift described above, LayV-F and other paramyxovirus species show differences in their surface-exposed residues in these regions that could define the unique antigenic properties of the different F proteins. These include a glycosylation site in LayV-F at N279 (Figure 4D), which is positioned between DI and DIII near several of the previously identified epitopes, specifically 15F, an HeV-neutralizing antibody and 5B3, a cross-reactive NiV/HeV-neutralizing antibody (30).

The sequences of both NiV-F and HeV-F have two plausible glycosylation sites at N64 and N67, both at the apex, though previous studies have established that only N67 is glycosylated (31, 32). The N65 glycosylation position in LayV-F is shifted by 8.2 Å away from the apex residues of HRA. This glycan was particularly visible in the cryo-EM reconstructions of the post-fusion conformation before local refinement, where the density at roughly the half-height point of the map corresponds to N65 (Supplemental Figure 3, lower right box). Interestingly, sequence alignment reveals that the LayV-F N65 glycosylation site is not homologous to either N64 or N67 of NiV/HeV, but rather is equivalent to residue 70 (Supplemental Figure 1). Viewing the fusion protein colored by sequence conservation further highlights the variability of this region (Figure 4E). While there are highly conserved residues at discrete points of the surface, namely the highly conserved disulfide bond near the apex, the surrounding region ranges from moderately conserved to highly variable, especially in surface-accessible regions of DIII and the HRA loop. This variability includes differing residue types between LayV and other henipaviruses, particularly at the apex site. The differences in the pre-fusion F protein trimer apex are highlighted by a negatively charged residue, D188, in both NiV and HeV, while the LayV-F protein instead has polar residues (N183, Q184) and an overall greater hydrophobic character. This residue arrangement in the pre-fusion LayV-F trimer apex is more similar to that of the PIV5 pre-fusion F protein (Figure 4F). This change in residue type is likely a source of antigenic variability between paramyxovirus F proteins. Consistent with this variability, antibody 12B2, that binds an epitope in pre-fusion NiV-F that includes the trimer apex (30, 33), did not show binding to LayV-F (supplemental Figure 2F). Therefore, despite similar F protein architecture, antigenic variability within paramyxoviruses is driven by variability of surface residues.

## Discussion

The continuing emergence of new paramyxovirus species necessitates heightened focus on a family that includes both some of the world’s deadliest pathogens in Nipah virus, and most infectious in measles and mumps. Members specifically of the *Henipavirus* genus have spread to humans through zoonotic transfer. While Nipah and Hendra viruses are spread to humans through bats, new henipaviruses have been detected in a range of species, such as shrews and rats. The role of animal reservoirs in the SARS-CoV-2 pandemic highlights the risk factor that a diverse range of animal hosts can be, further necessitating efforts to develop treatments or vaccines against henipaviruses. Structural determinations of *Henipavirus* glycoproteins serve as a foundation for immunogen or therapeutic design.

Though the structure of LayV-F is strikingly similar to other known *Henipavirus* F structures, key differences in surface characteristics at the apex have implications for pan-*Henipavirus* or pan-paramyxovirus vaccine development. When developing vaccinations for a virus that is quickly mutating, such as SARS-CoV-2, or for a virus with many distinct strains, as observed with henipaviruses, it is important to target relatively invariable sites for immunogen design. Our pre-fusion LayV-F protein structure helps to demonstrate the particularly variable nature of the apex and DIII in *Henipavirus* F proteins, a trait to be considered when selecting antigenic sites to target for a broad immunogenic response.

In contrast to existing structures of paramyxovirus fusion proteins that were determined in either pre- or post-fusion conformation, our study resolves the LayV-F pre- and post-fusion structures within the same cryo-EM dataset, suggesting that for LayV-F, being in solution at ambient temperatures may allow the proteins to sample enough conformational states such that some will lead to conversion. Though paramyxovirus fusion proteins must be cleaved to enable fusion peptide insertion, prior studies have observed uncleaved fusion proteins assuming a post-fusion conformation (34). Our structures reveal a highly stable configuration of the fusion peptide (FP) in the pre-fusion F conformation, buried within an interprotomer pocket, which must be disrupted to transition to the post-fusion conformation. These observations suggest that FP configuration is a source of metastability in the pre-fusion LayV-F protein, where a shift in the position of the FP or the residues surrounding it in the interprotomer pocket too far from its pre-fusion configuration may lead to irreversible conversion to the post-fusion form. This, together with the biochemical properties of the FP burial pocket, suggests tight control of FP positioning is an important part of pre-fusion metastability. While the use of an ectodomain construct without stabilizing mutations did allow for flexibility that hindered high resolution reconstruction, this made the relative lack of flexibility in the FP region more readily apparent, indicating a unique arrangement of FP conformation in paramyxoviruses as compared to other fusion proteins (Supplemental Figure 4, Supplemental Note 1).

An elusive question regarding paramyxovirus fusion is how the fusion process is first initiated. For the many virus families that utilize a fusion protein, the metastable pre-fusion state is maintained prior to a triggering event that initiates conformational changes. In many cases, this process involves cleavage and removal of entire domains of the fusion protein, such as with the removal of the S1 subunit from the SARS-CoV-2 spike protein prior to fusion. The paramyxovirus fusion proteins, however, have no such attachment subunit to remove. Instead, paramyxoviruses utilize a separate attachment protein, which is thought to pass on a triggering signal to the fusion protein after receptor binding (13, 14) (Figure 5A). For some paramyxovirus species, the attachment and fusion proteins have been shown to form complexes prior to receptor engagement (13) (Figure 5A). Structural characterization of such complexes had been elusive, but recently a structure of the human parainfluenza virus (hPIV3) attachment-fusion (HN-F) complex has been determined by cryo-electron tomography (35). One of the receptor-binding domains of HN is seen to bind at the apex of F, presumably stabilizing the pre-fusion conformation of F. While this might suggest a mechanism where the attachment protein functions as a clamp on the fusion protein, maintaining the pre-fusion state until receptor binding, studies assaying fusion competency have demonstrated that the fusion protein alone is not sufficient for virus and host cell fusion, which reinforces that paramyxovirus F proteins achieve inherent metastability that does not fully depend on an external clamp and indicates that fusion triggering involves a more complex mechanism of interaction between the two proteins (36, 37).

**Figure 5:**
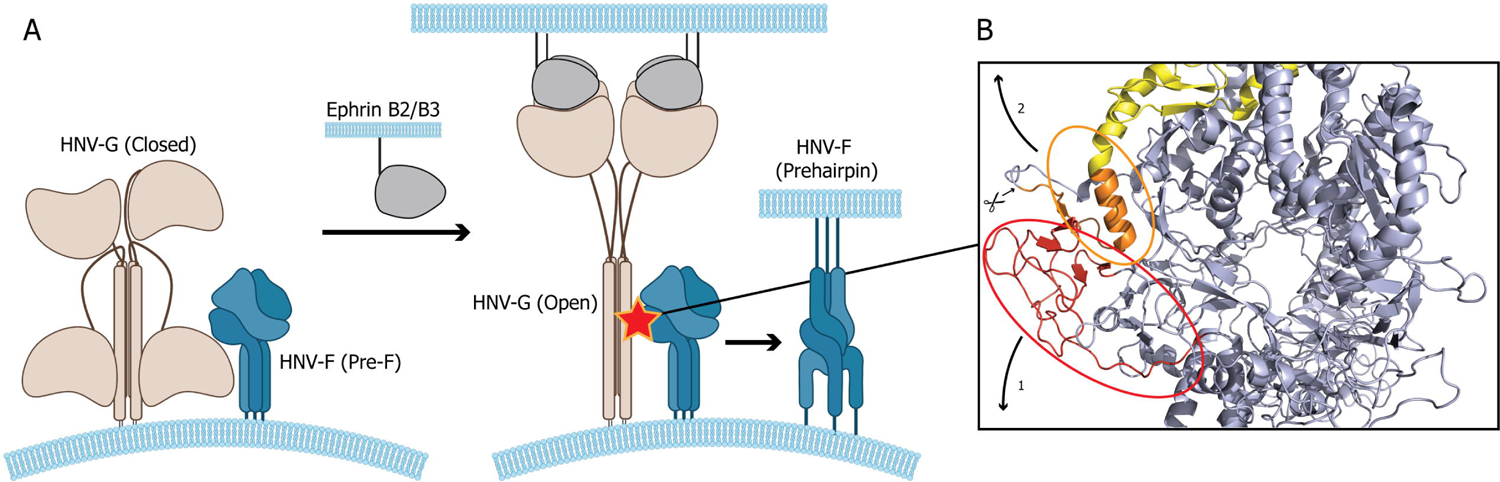
Fusion triggering hypothesis for paramyxoviruses. **A.** Existing model of *Henipavirus* (HNV) fusion protein triggering based on Liu *et al.* (2013) (36). HNV-G, brown, shown in a two-receptor binding domain down “closed” state, bound to HNV-F in pre-fusion conformation. Exposure to Ephrin B2/B3 receptor induces opening of HNV-G, triggering conversion of HNV-F from pre-fusion to the pre-hairpin state. The triggering interaction is labeled with a star. **B.** Hypothesized structural basis for the G-F interaction shown in **A**, wherein a two-step mechanism involving disruption of the fusion peptide burial pocket through action of the HNV-G stalk on DII frees the fusion peptide, allowing conformational conversion.

The structural similarities seen between the paramyxovirus fusion proteins of different species also extends to the receptor-binding head domains of paramyxovirus attachment proteins. In the hPIV3 HN-F structure, a loop from the HN head that inserts into a cavity formed at the trimer interface at the apex of F, which is noted to be one such area of structural conservation between different paramyxovirus genera, possibly revealing a conserved mode of complex formation (35). However, while the *Henipavirus* F proteins generally are generally glycosylated at their apexes, albeit if not at a strictly conserved residue, hPIV3 does not have a canonical N-X-S/T glycosylation motif at its apex and previous structures have not identified apex glycosylation for hPIV3F. The presence of a glycans in henipaviruses could add steric hinderance that may change the nature of this complex formation within this genus. It should be noted, however, that while hPIV3 has no canonical glycosylation motif at the apex, it does contain an N-X-C motif at residue 61 in the apex, which is a less frequently glycosylated sequence (38).

The fusion peptide in the pre-fusion LayV-F protein is in a configuration that is conserved among paramyxoviruses and distinct from other class-I fusion proteins (Supplemental Figure 4, Supplemental Note 1). It is not yet known which residues are involved in the interaction between the attachment and fusion proteins, but the critical role of the FP pocket created at the DII-DIII interface suggests that DII could be the site of the interaction. The triggering signal may be transferred by way of the attachment protein repositioning DII such that the fusion peptide is released from the pocket, irreversibly extending towards the apex (Figure 5B). This mechanism would be consistent with previous paramyxovirus studies that suggested an attachment protein-binding role for DII (39, 40). Further supporting this hypothesis is the likelihood that the FP in the pre-fusion conformation is in a high-energy state due to the kink in the helix comprised of the FP from T119 through HRA up to N148. Such a helix shape is often brought about by the presence of a proline at the kink, introducing bond-angle limitations that force this shape into the helix (41). However, in the helix containing the FP, no proline is present. The bent shape could be induced by the FP being buried within and locked into the hydrophobic FP pocket. When disrupted, the position of the FP would therefore move rapidly as the tension in the helix is resolved. Though it could be hypothesized that cleavage of the FP is the trigger that allows for rearrangement, our study, in line with others, has established that cleavage is not required for conformational conversion in paramyxovirus fusion proteins.

The existence of a high-energy state FP helix, held in place by an interprotomer DII, which is then disrupted by the action of an attachment protein offers a hypothesis for how *Henipavirus* attachment proteins activate their conjugate fusion proteins, and the basis for how paramyxovirus fusion proteins maintain their metastability in the pre-fusion state. Additional experiments, including biochemical studies assaying the effect of mutations in the FP-DII interface designed to stabilize or destabilize the interaction, will be needed to further explore this hypothesized mechanism of *Henipavirus* F metastability. Furthermore, while this study adds to the myriad of structural information on paramyxovirus F proteins, the breadth of information on paramyxovirus attachment protein structure remains comparatively lesser. Given the high variability in the stalk and neck domains of attachment proteins that may play a role in enabling complex formation and fusion activation, expanded structural information may be needed for effective experimental design that biochemically validates the nature of the attachment-fusion interaction.

## Materials and Methods

### Data Availability

Cryo-EM reconstructions and atomic models generated during this study are available at wwPDB and EMBD (https://www.rcsb.org; http://emsearch.rutgers.edu) under the accession codes PDB IDs 8FEJ and 8FEL and EMDB IDs EMD-29029 and EMD-29032.

### Plasmids

Gene synthesis for all plasmids generated by this study were performed and the sequence confirmed by GeneImmune Biotechnology (Rockville, MD). The fusion protein ectodomain constructs included F protein residues 1 to 438 (GenBank: UUV47205.1), a C-terminal T4 fibritin trimerization motif (FOLDON), a C-terminal HRV3C protease cleavage site, a TwinStrepTag and an 8XHisTag. The ectodomain was cloned into the mammalian expression vector pαH. Synthetic heavy and light chain variable domain genes for Fabs were cloned into a modified pVRC8400 expression vector, as previously described (5, 24, 25). Antibody variable regions for both heavy and light chain replaced the variable regions from the VRC01 antibody plasmid (42), with the IL-2 secretion signal and human kappa constant domain being used for light chain plasmids. All plasmids have been deposited to Addgene (https://www.addgene.org) under the codes 200390, 200391, 200392, 200398, and 200399.

### Cell culture and protein expression

For F ectodomains, GIBCO FreeStyle 293-F cells (embryonal, human kidney) were maintained at 37°C and 9% CO_2_ in a 75% humidified atmosphere in FreeStyle 293 Expression Medium (GIBCO). The Plasmid was transiently transfected using Turbo293 (SpeedBiosystems) and incubated at 37°C, 9% CO2, 75% humidity with agitation at 120 rpm for 6 days. On the day following transfection, HyClone CDM4HEK293 media (Cytiva, MA) was added to the cells. For antibodies, GIBCO Expi293F cells (embryonal, human kidney) were maintained at 37°C and 9% CO_2_ in a 75% humidified atmosphere in Expi293 Expression Medium (GIBCO). The Plasmid was transiently transfected using Expifectamine 293 (GIBCO) and incubated at 37°C, 9% CO2, 75% humidity with agitation at 120 rpm for 6 days. On the day following transfection, Expifectamine 293 Enhancers (GIBCO) were added to the cells.

### Protein purification

On the 6^th^ day post transfection, the fusion protein ectodomains were harvested from the concentrated supernatant, purified using StrepTactin resin (IBA LifeSciences) and size exclusion chromatography (SEC) using a Superose 6 10/300 GL Increase column (Cytiva, MA) equilibrated in PBS buffer (Thermo Scientific 137mM NaCl, 2.7mM KCl, pH 7.4, 10 mM Phosphate Buffer, 1.8mM Potassium Phosphate Monobasic). Antibodies were purified using Protein A affinity (Thermo Scientific) and SEC using a Superose 6 10/300 GL Increase column (Cytiva, MA) equilibrated in PBS buffer (Thermo Scientific 137mM NaCl, 2.7mM KCl, pH 7.4, 10 mM Phosphate Buffer, 1.8mM Potassium Phosphate Monobasic). All steps of the purification were performed at room temperature and in a single day. Protein quality was assessed by SDS-Page using NuPage 4-12% (Invitrogen, CA). The purified proteins were flash frozen and stored at −80 °C in single-use aliquots. Each aliquot was thawed by a 5-minute incubation at 37 °C before use.

### Cryo-EM

Purified ectodomain was diluted to a concentration of 1.5 mg/mL in PBS (described above) with 0.0005% DDS and 0.38% glycerol added. A 2.2µL drop of protein was deposited on a Quantifoil R1.2/1.3 grid (Electron Microscopy Sciences, PA) that had been glow discharged for 10 seconds using a PELCO easiGlow™ Glow Discharge Cleaning System. After a 30-second incubation in >95% humidity, excess protein was blotted away for 2.5 seconds before being plunge frozen into liquid ethane using a Leica EM GP2 plunge freezer (Leica Microsystems). Frozen grids were imaged using a Titan Krios (Thermo Fisher) equipped with a K3 detector (Gatan). Movie frame alignment was carried out using UNBLUR (43).The cryoSPARC (44) software was used for data processing. Phenix (45), PyMOL (46), ChimeraX (47) and Isolde (48) were used for model building and refinement. Phenix was first used to fit the initial models into maps. ISOLDE was used to manually adjust residues to address rotamer and Ramachandran outliers. Phenix was again used to perform real-space refinement, energy-minimization of side-chain positions, and to set B-factors during an ADP-only real-space refinement. The resolution of both structures did not allow for experimental determination of side chain positions, therefore energy minimization followed by optimization of rotamer positions described above are the basis for side chains in the model.

### SDS-PAGE

Prepared 1, 3, or 8µg of sample with Laemmli sample buffer (BioRad), PBS buffer, and with or without 300mM DTT (Reduced/Non-Reduced). Loaded to NuPAGE 4-12% Bis-Tris gel (Invitrogen) and ran at 175V with MES-SDS running buffer until complete. Stained with Coomassie blue (Novex) for 30 minutes before water destain and imaging.

### Differential scanning fluorimetry

DSF assay was performed using Tycho NT. 6 (NanoTemper Technologies). Spike ectodomains were diluted to approximately 0.15LJmg/mL. Intrinsic fluorescence was measured at 330 nm and 350 nm while the sample was heated from 35 to 95 °C at a rate of 30°C/min. The ratio of fluorescence (350/330 nm) and inflection temperatures (Ti) were calculated by the Tycho NT. 6 apparatus.

### Difference distance matrices (DDM)

DDM plots were generated using the Bio3D package (Grant *et al*., 2021) implemented in R (R Core Team, 2018. R: A language and environment for statistical computing. R Foundation for Statistical Computing, Vienna, Austria. http://www.R-project.org/).

### Bio-layer Interferometry (BLI)

Antibody binding to *Henipavirus* F proteins was assessed using BLI on an Octet RED 384 (Sartorius, formerly ForteBio) with Forte Bio Kinetics Buffer. All binding assays were performed at 30°C. Antibodies were immobilized on Anti-Human Fc Capture tips (ForteBio), loaded at 20µg/mL for 300s. F proteins were used as analytes at 100nM, with an association time of 480s and dissociation time of 600s. Sensorgram data were reference-subtracted and analyzed using the Octet Analysis Studio software (Sartorius), with a reference tip for each immobilized antibody and reference sample for each analyte.

### Sequence Alignment

Paramyxovirus F sequences aligned with Clustal Omega tool (49). Output alignment was colored and assigned consensus residue types by MVeiw (50). Sequences obtained from either Uniprot or Genbank. Uniprot accession numbers: NiV-F: Q9IH63, HeV-F: O89342, hPIV3-F: P06828, PIV5-F: P04849, MeV-F (ICB Strain): Q786F3, MojV-F: W8SKT3, CedV-F: J7GX38, hPIV1-F: P12605, NDV-F (Texas Strain): P26628, SeV-F (Z Strain): P04855. Genbank accession numbers: LayV-F: UUV47205.1, GakV-F: QYO90524.1, DarV-F: QYO90531.1.

## Acknowledgments

Figure 5 was made using resources from BioRender. Cryo-EM data were collected at the Duke Krios at the Duke University Shared Materials Instrumentation Facility (SMIF), a member of the North Carolina Research Triangle Nanotechnology Network (RTNN), which is supported by the National Science Foundation (award number ECCS-2025064) as part of the National Nanotechnology Coordinated Infrastructure (NNCI). This study utilized the computational resources offered by Duke Research Computing (http://rc.duke.edu; NIH 1S10OD018164-01) at Duke University.

## List of figures

**Supplemental Figure 1:** Sequence Alignment of *Paramyxoviridae* Fusion Proteins. Color-coded sequence alignment of select *Paramyxoviridae* Fusion proteins compared to the LayV-F protein.

**Supplemental Figure 2:** Purification and Antibody Binding of *Henipavirus* Proteins. Purification and quality control results for several *Henipavirus* fusion protein constructs with BLI binding data of anti-NiV antibodies against these constructs.

**Supplemental Figure 3:** Cryo-EM processing of LayV-F structures. Cryo-EM processing of LayV-F in pre- and post-fusion conformation, showing representative micrographs, 2D classes, intermediate 3D refinements, and workflow.

**Supplemental Figure 4:** Fusion Peptide comparison. Comparison of fusion peptide positions across several notable virus species.

**Supplemental Figure 5:** Fusion Peptide Pocket Enlarged view of a panel from figure 3 depicting the fusion peptide burial pocket.

## Notes

### Competing Interest Statement

The authors have declared no competing interest.

### Summary of Updates

Discussion section now comments on a relevant paper to paramyxovirus attachment-fusion protein complexes. A supplemental figure has been replaced with a different figure.

