## Supplemental Figures 1-5 for "Structures of Langya virus fusion protein ectodomain in pre and post fusion conformation"

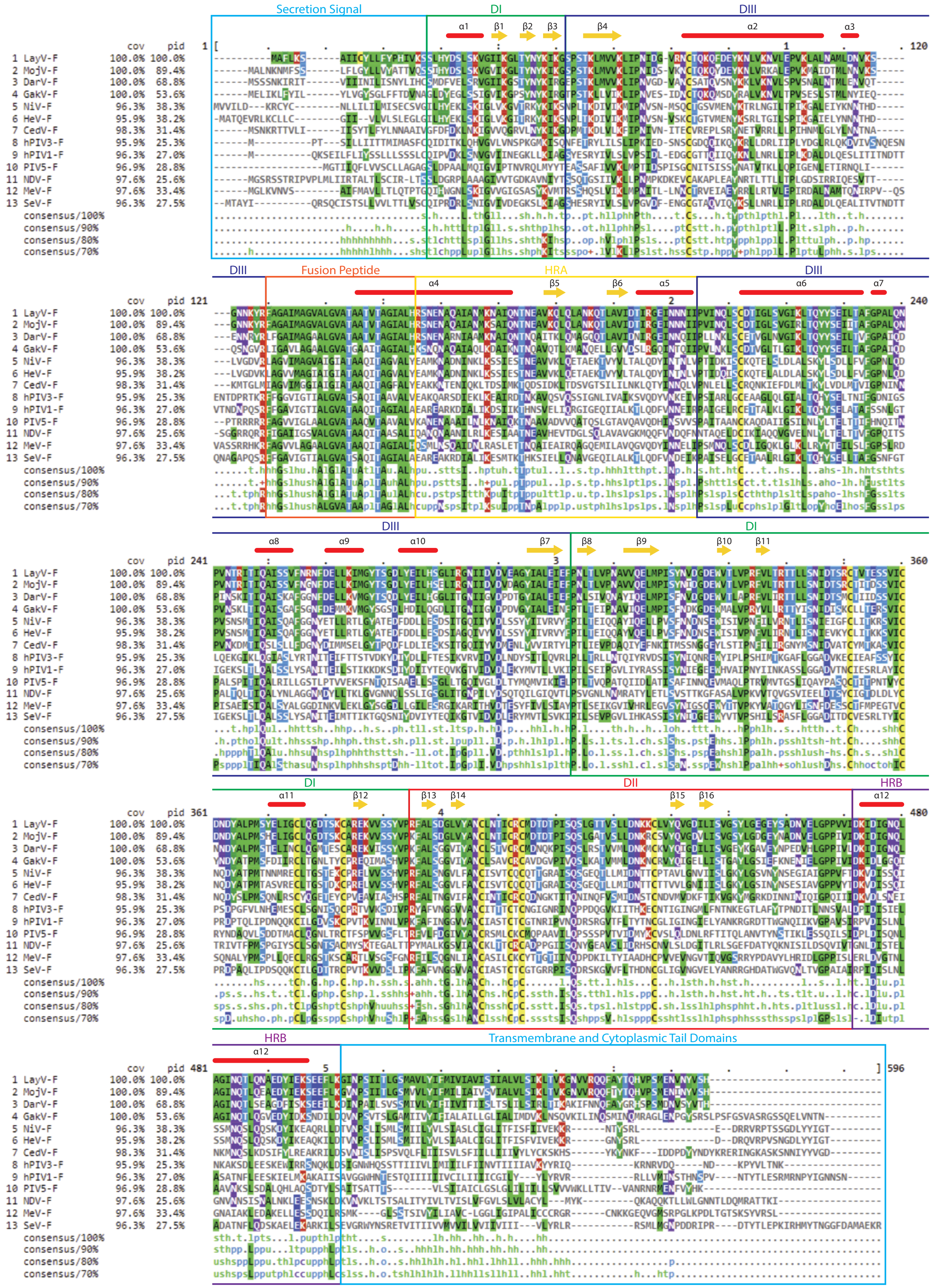


**Supplemental Figure 1: Sequence Alignment of *Paramyxoviridae* Fusion Proteins.**

Sequence alignment of select paramyxovirus fusion proteins with LayV-F used as the reference sequence. Cov: percent sequence coverage. Pid: percent sequence identity. Residues are colored by type, per MView standard (50). Blue: Alcohol, Green: Hydrophobic, Dark Blue: Negative charge, Red: Positive charge, Purple: Polar, Yellow: Cysteine. Consensus residue types assigned per MView standard. Alcohol: (o), Aliphatic: (l), Aromatic: (a), Charged: (c), Hydrophobic: (h), Negative: (-), Polar: (p), Positive: (+), Small: (s), Tiny: (u), Turnlike: (t).

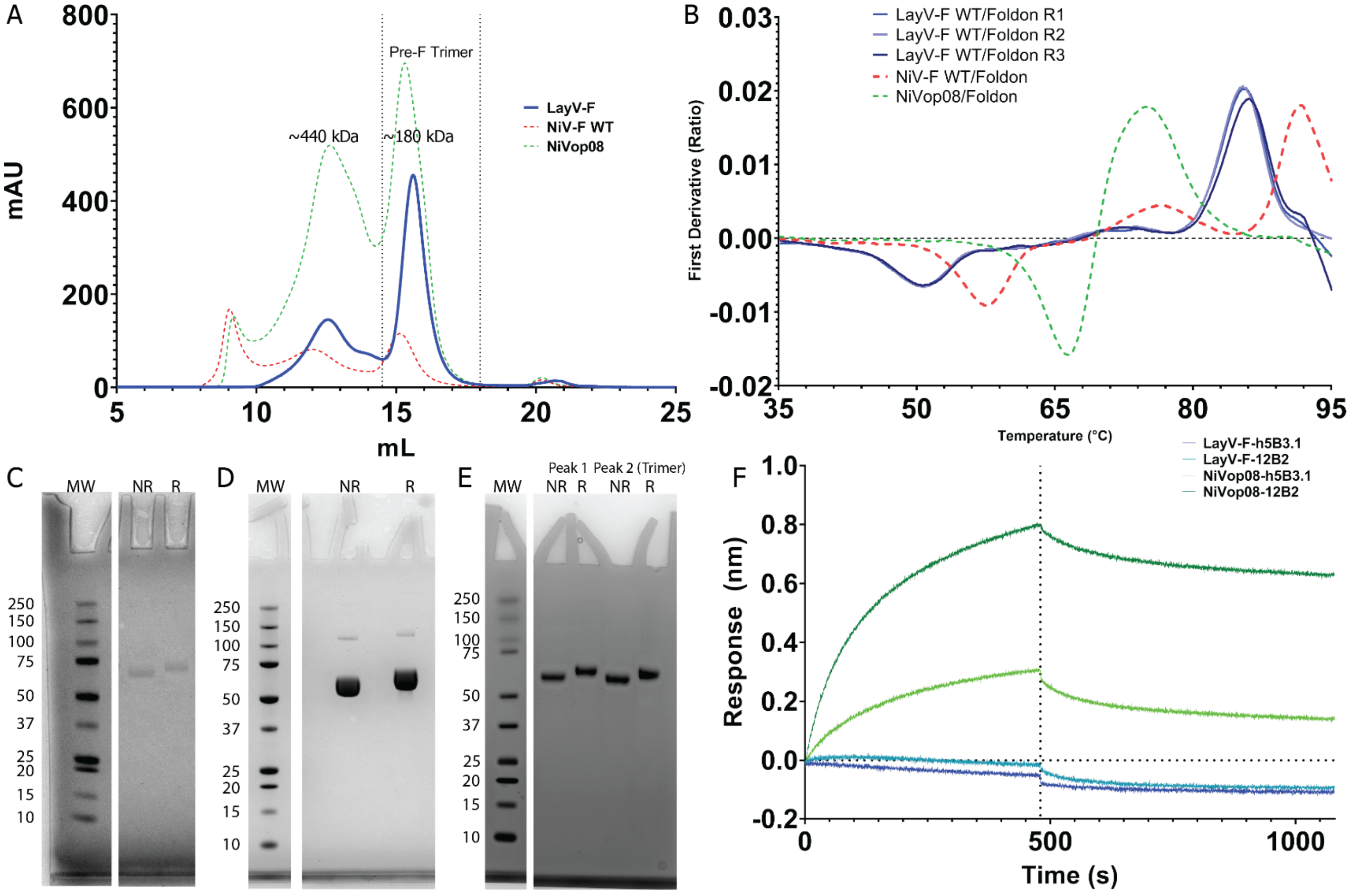


**Supplemental Figure 2: Purification and Antibody Binding of *Henipavirus* Proteins.**

**A.** Size-exclusion chromatogram of *Henipavirus* fusion protein purification. LayV-F, blue, NiV-F WT, dashed red, pre-fusion stabilized NiVop08 (16), dashed green. The fractions identified as the ectodomain trimer at 180kDa are indicated with a vertical dashed line **B.** Differential scanning fluorimetry profiles of the products from A, LayV-F measured in triplicate. **C-E.** SDS-PAGE of *Henipavirus* constructs from A. Molecular weight markers, MW, non-reducing conditions, NR, reducing conditions, R., **C.** LayV-F loaded at 1μg per lane. **D.** LayV-F loaded at 8μg per lane. **E.** NiVop08 loaded at 3μg per lane, with left two bands sampling the first peak at 440kDa and the second two bands sampling the trimer peak at 180kDa. **F.** BLI sensorgram of anti-NiV-F antibodies h5B3.1 and 12B2 binding to immobilized LayV-F or NiVop08 (30, 33).

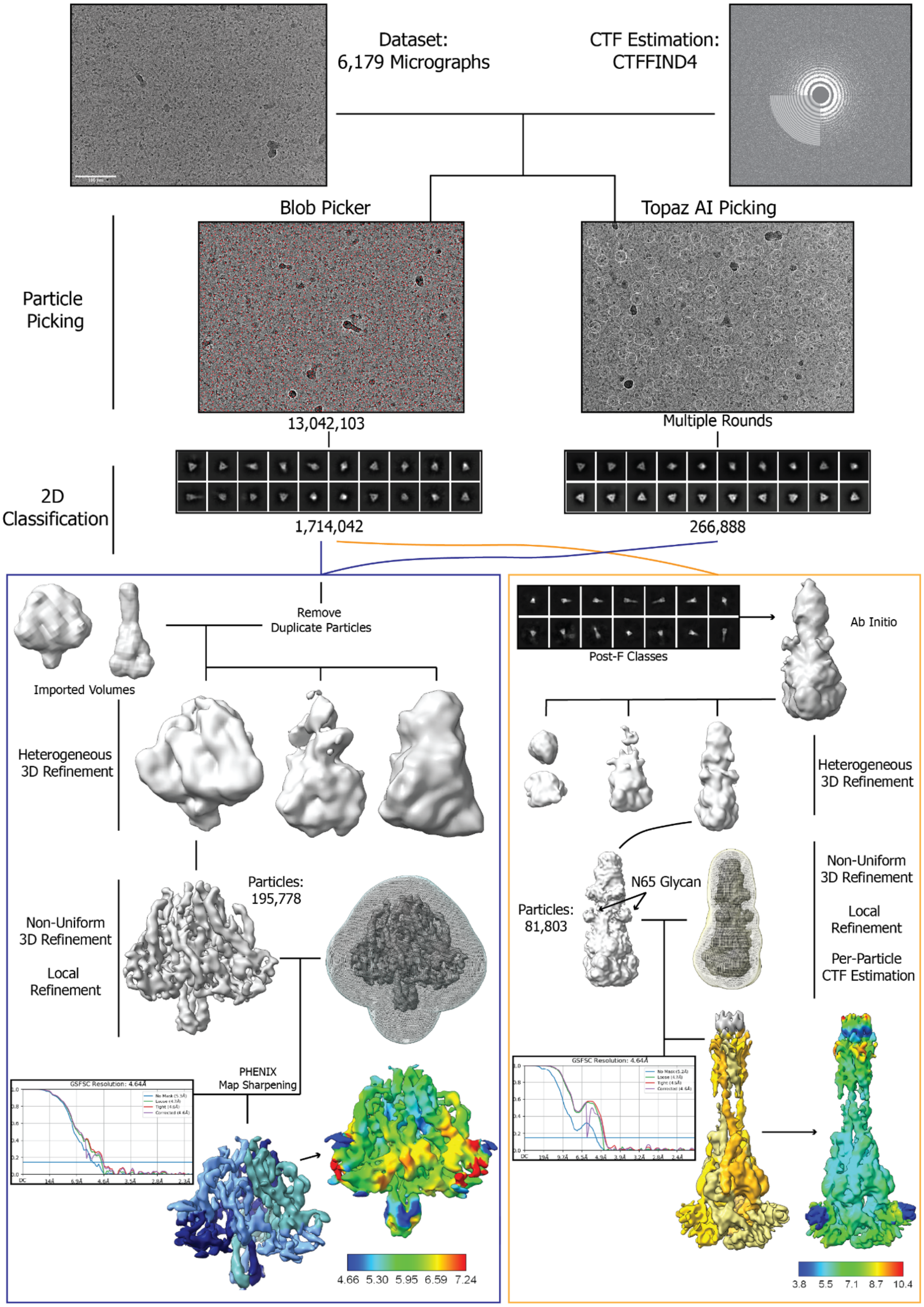


**Supplemental Figure 3: Cryo-EM processing of LayV-F structures.**

Cryo-EM processing of pre- and post-fusion LayV-F structures. Steps performed using CryoSPARC (44, 53). CTF Estimation performed on micrographs followed by default blob picker algorithm and a trained AI-based algorithm (Topaz) for particle picking. Particles for the post-fusion state were obtained through the blob picker, while particles contributing to the pre-fusion state came from both sets. Heterogeneous refinement was used to classify and further sort particles. After non-uniform refinement, a local refinement was performed on the entire structure. The supplemental map of the post-fusion state prior to local refinement where the N65 glycan is detected has also been deposited.

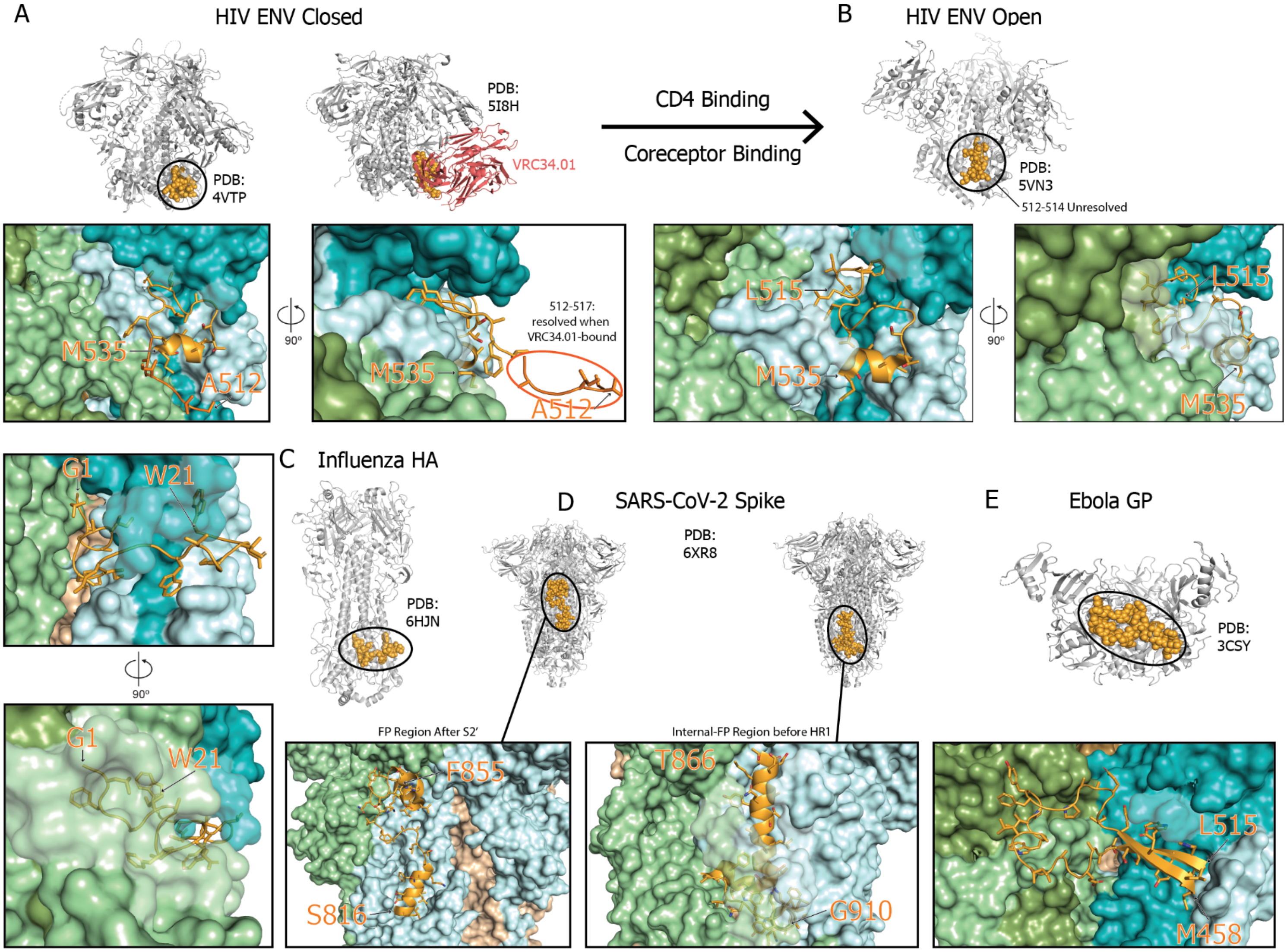


**Supplemental Figure 4: Fusion Peptide comparison.**

**A-E:** Comparison of fusion peptide location in other viral fusion proteins. In the top portion of each panel, fusion proteins are shown in cartoon representation in gray, with the fusion peptide as orange spheres, and in the bottom portion, fusion peptides are shown as cartoon representation with side chains as sticks, orange, with the surface of the rest of the fusion protein colored based on protomer. The most proximal protomer to the highlighted fusion peptide is colored cyan, with the neighboring protomers colored green and tan. In cases where there are are multiple subunits in the fusion protein, the attachment subunit is colored a darker shade than the corresponding fusion subunit. **A.** HIV Envelope (Env) closed conformation, with (PDB: 4TVP) and without fusion peptide-targeting antibody VRC34.01 (PDB: 5I8H), shown in cartoon, red. **B**. HIV Env in open conformation (PDB: 5VN3). **C.** Influenza HA in closed conformation (PDB: 6HJN). **D.** SARS-CoV-2 Spike in pre-fusion, receptor-binding domain down conformation (PDB: 6XR8). **E.** Ebola GP in pre-fusion conformation (PDB: 3CSY). (54-59)

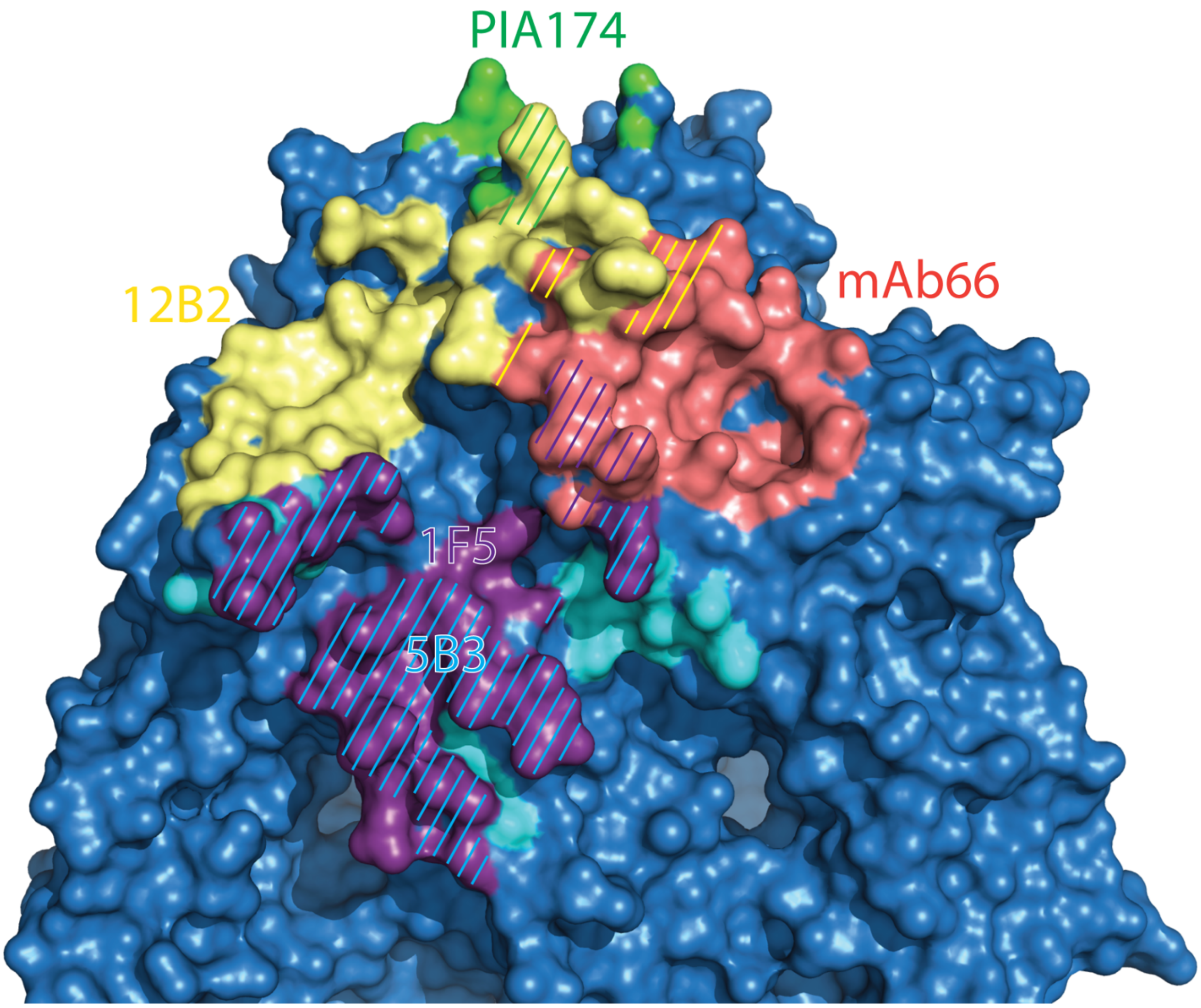


**Supplemental Figure 5: Paramyxovirus F Antibody Binding Footprints**

Binding footprints of anti-F antibodies shown in figure 4C. NiV-F from PDB 7KI4 is shown both in surface view. Sites interacting with antibodies are colored per the color scheme from 4C. Areas with overlap are represented with hatched lines of the appropriate colors.
