## Supplemental Note 1, Supplemental Table 1 for "Structures of Langya virus fusion protein ectodomain in pre and post fusion conformation"

Comparison of the LayV-F fusion peptide (FP) arrangement to other viral fusion peptides highlights the unique nature of the paramyxovirus FP. Whereas a large portion of the LayV-F FP is buried within an interprotomer pocket, the HIV envelope protein (Env), the FP remains largely surface exposed in the closed conformation (Supplemental Figure 4A) (59). The most N-terminal residues of the HIV-1 FP are often unresolved in structures, further highlighting the flexibility of the FP in this Env conformation. During receptor-induced Env opening, the FP begins to be pulled into the core of the protein, and though not buried, it has a somewhat lesser degree of surface exposure (Supplemental Figure 4B) (58). In contrast, the Influenza Hemagglutinin protein (HA), in the pre-fusion state, maintains the N-terminal end of the FP in a hydrophobic cavity at the core of the protein (Supplemental Figure 4C) (56). While the Influenza HA FP is not mobile and flexible as with the HIV-1 FP in the closed state of Env, the Influenza HA FP arrangement does not constitute as true burial, as the FP is inserted into a recess in the structure but not occluded by other residues. The exact nature of the SARS-CoV-2 spike FP is uncertain, as several regions in S2 may play a role. Many structures have identified the stretch of residues directly following the S2’ cleavage site as the fusion peptide. This region is not buried, and like the FP of HIV Env, is often poorly resolved in spike structures (Supplemental Figure 4D) (55). However, a recent study (60) indicates that it is residues 866-910 that insert into the host membrane. Close examination of the SARS-CoV-2 sequence reveals that the residues that in the middle of this FP region, which form an internal FP, much more closely matching the glycine and alanine-rich sequences of other known FPs than those directly following the S2’ site. This FP arrangement has a much greater degree of similarity in pre-fusion arrangement to paramyxoviruses. The FP region is almost completely buried by the surrounding domain and has limited surface accessibility (Supplemental Figure 4D). While this FP burial is more comparable to paramyxoviruses, the usage of an internal FP loop draws a comparison to the Ebola glycoprotein (GP) FP, also an internal FP. This Ebola FP region is not buried in the GP structure, instead forming the exterior of the fusion subunit. Though it is an internal FP, as with SARS-CoV-2, the fusion loop is positioned between two beta strands rather than two alpha helices (Supplemental Figure 4E) (54).

| **Table S1. Cryo-EM data collection and refinement statistics.** | | |
| --- | --- | --- |
|  | **LayV-F Pre-Fusion** | **LayV-F Post-Fusion** |
| **PDB ID** | 8FEJ | 8FEL |
| **EMDB ID** | EMD-29029 | EMD-29032 |
| **Data Collection and processing** | |  |
| Microscope | FEI Titan Krios | |
| Detector | Gatan K3 | |
| Magnification | 81000 | |
| Voltage (kV) | 300 | |
| Electron exposure (e-/Å^2) | 56.3 | |
| Defocus Range (µm) | 2 to 0.8 | |
| Pixel size (Å) | 1.08 | |
| Reconstruction software | cryoSPARC | |
| Symmetry imposed | C3 | |
| Initial particle images (no.) | 13,042,103 | |
| Final particle images (no.) | 195,778 | 81,802 |
| Map resolution (Å) | 4.64 | 4.64 |
| FSC threshold | 0.143 | 0.143 |
| **Model composition** |  |  |
| Nonhydrogen atoms | 10467 | 9204 |
| Protein residues | 1368 | 1191 |
| **R.M.S. deviations** |  |  |
| Bond lengths (Å) | 0.003 | 0.012 |
| Bond angles (°) | 0.864 | 1.698 |
| **Validation** |  |  |
| MolProbity score | 2.01 | 1.62 |
| Clashscore | 6.26 | 3.13 |
| Poor rotamers (%) | 4.3 | 1.04 |
| **Ramachandran plot** |  |  |
| Favored regions (%) | 96.92 | 91.56 |
| Allowed (%) | 2.86 | 8.44 |
| Disallowed regions (%) | 0.22 | 0 |
